# Nucleoporin 133 deficiency leads to glomerular damage in zebrafish

**DOI:** 10.1101/352971

**Authors:** Chiara Cianciolo Cosentino, Alessandro Berto, Michelle Hari, Johannes Loffing, Stephan C. F. Neuhauss, Valérie Doye

## Abstract

Although structural nuclear pore proteins (nucleoporins) are seemingly required in every cell type to assemble a functional nuclear transport machinery, mutations or deregulation of a subset of them have been associated with specific human hereditary diseases. In particular, previous genetic studies of patients with nephrotic syndrome identified mutations in *Nup107* that impaired the expression or the localization of its direct partner at nuclear pores, Nup133. In the present study, we characterized the zebrafish *nup133* orthologous gene and its expression pattern during larval development. Morpholino-mediated gene knockdown revealed that Nup133 depletion in zebrafish larvae leads to the formation of kidney cysts, a phenotype that can be rescued by co-injection of wild type mRNA. Analysis of different markers for tubular and glomerular development shows that the overall kidney development is not affected by *nup133* knockdown. On the other hand, we demonstrate that *nup133* is essential for the organization and functional integrity of the pronephric glomerular filtration barrier, as its downregulation results in proteinuria and moderate foot process effacement, mimicking some of the abnormalities typically featured by patients with nephrotic syndrome. These data indicate that *nup133* is a new gene required for proper glomerular structure and function in zebrafish.

## Introduction

Efficient and regulated bidirectional transport between the cytoplasm and the nucleus is an essential process in all eukaryotic cells. This function is achieved by nuclear pore complexes (NPCs), huge assemblies anchored within the nuclear envelope and composed of about 30 different proteins, termed nucleoporins (Nups) (reviewed in ^1^). Despite the universal role of NPCs in all nucleated cells, some Nups are linked to human hereditary diseases affecting specific cell types or organs (reviewed in ^2,3,4^).

In particular, genetic studies have implicated a restricted number of structural nucleoporins in specific kidney diseases termed nephrotic syndromes (NS). NS arise from defects or damages that impair the selectivity of the glomerular filtration barrier and lead to massive proteinuria and hypoalbuminemia, which in turn cause edema and dyslipidemia. The glomerular filtration barrier (GFB) surrounds the glomerular capillaries and comprises three layers: (i) a fenestrated endothelium, (ii) a basement membrane, and (iii) the podocytes. The latter are highly specialized epithelial cells characterized by long and thin cytoplasmic projections, termed foot processes (FPs), that interdigitate and are connected by specialized cell-cell junctions, the slit diaphragms (reviewed in ^5^). While most patients with childhood-onset NS respond well to steroid treatments, 10-20% of the affected children do not achieve remission upon corticosteroid therapy. Steroid-resistant NS (SRNS) is associated with a high risk of progression to end-stage renal disease (ESRD) ^6^. It frequently manifests histologically as focal segmental glomerulosclerosis (FSGS), characterized by scattered scarring of some glomeruli and is often associated with retractions (“effacement”) of podocytes foot processes (reviewed in ^7^).

Although nonhereditary forms of SRNS seem to be prevalent, studies over the last years have identified over 50 dominant or recessive single-gene mutations in a significant percentage (30%) of patients with early-onset SRNS and FSGS (reviewed or discussed in ^6,8–11^). While some of these genes have podocyte-specific or -restricted functions, these studies also unveiled the implication of multiple cellular processes in the establishment or maintenance of the glomerular filtration barrier ^7,12,13^. In particular, these genetic studies have identified mutations in Nup93 and Nup205, two constituents of the inner ring of the NPC ^14^ and in Nup107, a constituent of the Y-complex (Nup107/160-complex) that builds up the cytoplasmic and nuclear rings of the NPC ^15,16,17^. Mutations within *Nup107* were also identified in patients with a rare co-occurrence of microcephaly with nephrotic syndrome, similar to Galloway-Mowat syndrome (GAMOS) ^18^. While patients with GAMOS-like presentation had a strong reduction in Nup107 protein level accompanied by decreased levels of Nup133, its direct partners within the Y-complex ^18^, another SRNS-linked mutation affecting Nup107 was shown to impair its interaction with Nup133 ^16^. These data thus pointed towards a possible implication of Nup133 in NS.

In mice, a previous characterization of a *Nup133* null (*mermaid, or merm*) mutant revealed that mouse embryos lacking a functional *Nup133* allele developed through midgestation but die at e9.5-e10.5 ^19^. While this indicates that Nup133 is not an obligate NPC component, *merm* mutants displayed abnormalities in a number of tissues, indicating that cell differentiation towards several epiblast-derived lineages likely requires Nup133 ^19^. However, the possible contribution of Nup133 to kidney development or function has never been assessed. To address this question, we used morpholino-mediated *nup133* inactivation in zebrafish (*Danio rerio, Dr*), a well-established vertebrate model to study kidney development and model renal diseases ^20–23^. We report here that knockdown of zebrafish *nup133*, while not impairing early stages of kidney development, leads to glomerular abnormality that mimic nephrotic syndrome.

## Results

### Zebrafish *nup133* ortholog is broadly expressed at early stages and becomes restricted to specific tissues at later stages

Query of the latest version of the genome databases identified a provisional zebrafish *Dr nup133* gene (<u>ZFIN:ZDB-GENE-040426-2941</u>; Ensembl:ENSDARG00000010078) containing 26 exons located on the forward strand of chromosome 1. The open reading frame is predicted to encode a protein of 1136 amino acids (aa) that shares 62.3% overall amino acid identity with human Nup133 (Supplementary Fig. S1 online).

To determine the spatio-temporal localization of *nup133* transcripts in zebrafish embryonic tissues, we performed whole-mount in situ hybridization (ISH) at different developmental stages (Fig. 1a-f). The *nup133* sense RNA probe was used as negative control (Supplementary Fig. S2 online). Ubiquitous expression of *nup133* was observed at shield stage (5 hours post fertilization, hpf, Fig. 1a and Supplementary Fig. S2 online). By 24 hpf, *nup133* mRNA was detected in the central nervous system, with higher levels of expression in the retina, the tectum and the cerebellum (Fig. 1b). At 3 days post fertilization (dpf), in addition to a diffuse staining notably in the brain, we found evidence of *nup133* mRNA enrichment in the liver, in the intestine and in neuromasts of the lateral line organ (Fig. 1c,d). At 5 dpf, the overall expression of *nup133* is weaker, but the mRNA is still visible in the brain, and enriched in the liver, the intestine and the swim bladder (Fig. 1e,f). Cross sections at 4 and 5 dpf confirmed the enrichment of *nup133* mRNA in the liver and highlighted the presence of *nup133* mRNA also in the pronephric proximal tubules and the glomerulus (Fig. 1g,h). Because structural nucleoporins were reported to be long-lived proteins, with a very low turnover in at least some non-dividing cells ^24,25^, we next sought to determine whether higher *nup133* mRNA expression in the larvae was associated with highly proliferating tissues. To evaluate cell proliferation, we treated 5 dpf zebrafish larvae with the thymidine analog 5-ethynyl-2′-deoxyuridine (EdU) for 1 h and assayed for EdU-positive cells in the different tissues of the larvae. As shown in Figure 1i, EdU positive cells are more frequent in the developing fins, the intestine and the liver, that with exception to the liver, do not correspond to tissues with major enrichment of *nup133* mRNA. This indicates that *nup133* expression in zebrafish is not directly correlated with cell proliferation.

**Figure 1.**
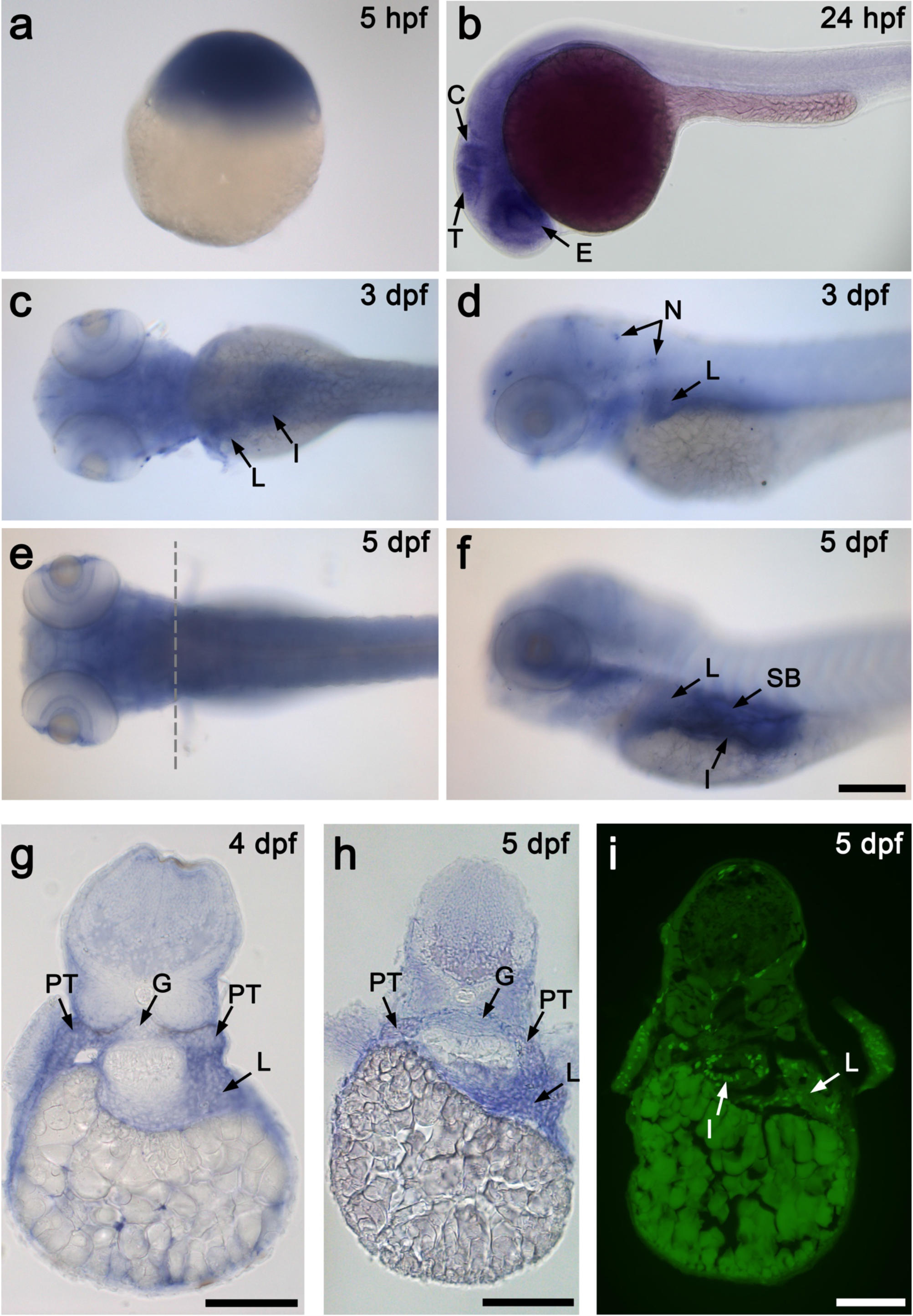
Expression of *nup133* in the developing zebrafish detected by *in situ* hybridization (ISH). (**A**) Whole mount ISH with *nup133* antisense probe of embryos at: (**a**) sphere stage (5 hpf; embryo shown with animal pole to the top); (**b**) 24 hfp (lateral view); (**c-f**) 3 and 5 dpf (left panels: dorsal view; right panels: lateral view). Arrows point to tissues with enriched expression of *nup133*. Abbreviations: E: eyes; T: tectum; C: cerebellum; L: liver; G: glomerulus; I: intestine; N: neuromasts; SB: swim bladder; PT: proximal tubules. Scale bars, 200 μm. (**B**) (**a, b**) Transverse sections of 4 and 5 dpf embryos at the level of the pectoral fins (as shown in the dotted line in **A, e**) confirm *nup133* expression in the liver, and show in addition a diffuse staining in the proximal tubules and the glomerulus. (**c**) Transverse crysections in the same region of 5 dpf larvae incubated with EdU for 1h prior to fixation and EdU detection. Scale bars, 100 μm.

### *nup133* knockdown causes glomerular cysts in zebrafish

In order to characterize the *in vivo* function of *nup133* in zebrafish larvae, we generated knockdown larvae using splice-blocking antisense morpholino oligos (MO) targeting the exon-intron boundary (splice donor E3I3) of exon 3 and the intron-exon boundary of exon 4 (splice acceptor I3E4) of the *nup133* gene (Fig. 2a). Retention of intron 3 in *nup133* transcripts is predicted to produce a truncated protein of 124 aa. Reverse transcription-PCR (RT-PCR) demonstrated that the morpholinos interfere with the splicing of exon 3, as revealed by the sequencing of the additional RT-PCR product detected in MO-treated compared to control embryos (Fig. 2b and Supplementary Fig. S3 online). Following *nup133* MOs injections, zebrafish larvae frequently developed pericardial edema and exhibited an expansion of the glomerulus detectable at 3 dpf by the formation of pronephric cysts (Fig. 2c and Supplementary Fig. S3 online). Using the *Tg(wt1b:eGFP)* transgenic line ^26^, in which podocytes and proximal pronephric tubules express EGFP under the *wt1b* promoter, the glomerular expansion could be directly observed under a fluorescence microscope (Fig. 2d, arrows). Analysis of semi-thin transverse sections of the glomerulus and proximal tubules at 5 dpf confirmed that the Bowman’s space of the glomerulus was dilated in *nup133* morphants compared with the control larvae (Fig. 2f). In contrast, no major dilatation of the proximal tubules was observed (Fig. 2f).

**Figure 2.**
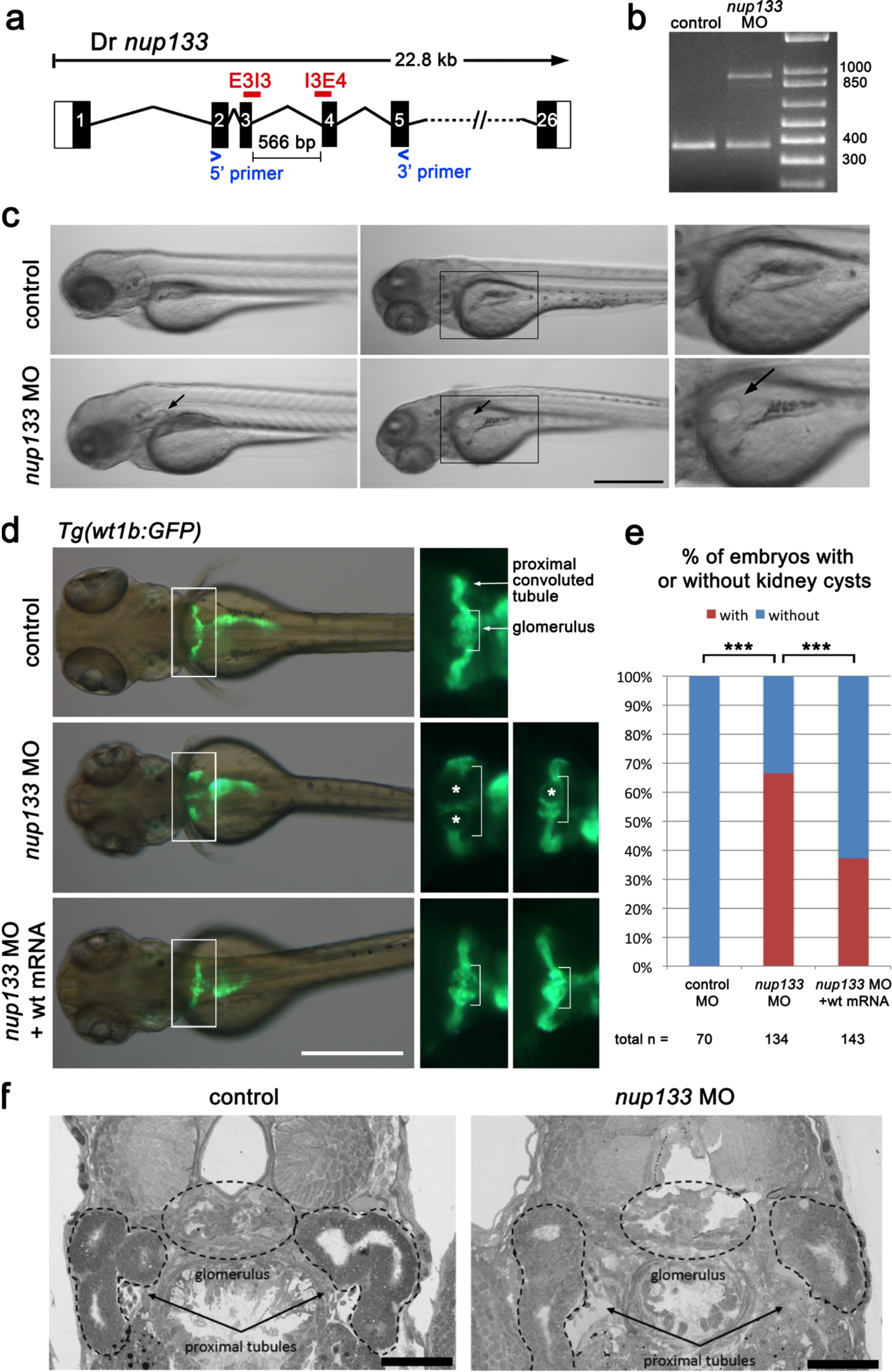
Morpholino (MO) knockdown of *nup133* causes glomerular expansion in zebrafish. (**A**) Exon structure of *Danio rerio* (*Dr*) *nup133* around the binding sites of the E3I3 and I3E4 splice morpholinos. Blue arrowheads indicate the position of the RT-PCR primers. The size of intron 3 is indicated. (**B**) RT-PCR from lysates of 48 hpf uninjected embryos (control) and of embryos coinjected with the two splicing morpholinos (*nup133* MO) reveal an additional band caused by retention of intron 3. (**C**) Gross morphology of 3 dpf control and *nup133* MO embryos (left panels: lateral view; middle panel: dorso-lateral view from two distinct embryos). Scale bar, 500 μm. Two-fold magnification of the indicated area is shown in the rightmost panels. Arrows indicate the pronephric cysts detected in the *nup133* MO embryos. (**D**) Dorsal view of 3dpf *Tg(wt1b:EGFP)* embryos uninjected (control, top panels), injected with *nup133* MO (middle panels), or sequentially injected with *3xHA-mCherry-Dr nup133* mRNA (wt mRNA) and *nup133* MO (bottom panels). Overlays of transmission (gray) and GFP-signal (green) images reveal the glomerulus, proximal tubules, and exocrine pancreas. Scale bar, 500 μm. Two-fold magnification of the indicated area and of the same area from a distinct larvae are shown in the right and rightmost panels, respectively. The glomerular structure is indicated in brackets. Note that the cystic dilations of the pronephros in *nup133* MO (middle panels, asterisks). (**E**) Relative proportion of embryos showing or not kidney cysts at 3 dpf. For each condition, the total number of embryos analyzed is indicated (n=, quantified in 2 distinct experiments for control MO injections and 5 experiments for *nup133* MO injections. See also Supplementary Fig. S4 online). Unlike the embryos injected with control MO, those injected with *nup133* MO frequently feature kidney cysts. On the other hand *nup133* MO + *wt* mRNA showed significantly fewer cysts than *nup133* MO alone. *** P< 0.0001 using a Fisher exact probability test. (**F**) Toluidine blue stained sections of 5 dpf uninjected control (left panel) and *nup133* MO injected larvae (right panel) showing the glomerulus and the proximal tubules (indicated with dotted lines). Note the cystic dilation of the glomerulus in the *nup133* MO larvae. Scale bars 50 μm.

To establish specificity of MOs effects, we determined whether *nup133* MO phenotypes could be rescued by co-injection of a synthetic zebrafish *nup133* (*Dr nup133*) mRNA. The *Dr nup133* mRNA was fused to a triple-hemagglutinin (HA) epitope and mCherry that enabled us to confirm the expression of the resulting 3xHA-mCherry-Dr Nup133 protein by western blot (Supplementary Fig. S4 online). Co-injection of the splice MOs with the *3xHA-mCherry-Dr nup133* mRNAs reduced significantly the percentage of larvae with glomerular cysts (Fig. 2e and Supplementary Fig. S4 online). This demonstrates that the glomerular phenotype observed in *nup133* morphants is due to a specific effect of *nup133* knockdown.

### *nup133* morphants properly express molecular markers of kidney development

We next wanted to determine whether the glomerular expansion in *nup133* morphants is caused by a developmental defect of the glomerulus and/or of the pronephros. For this purpose, we used whole-mount ISH to screen established markers of pronephros and glomerulus development. We first assayed expression of the intermediate mesoderm marker *pax2a* (*paired box gene 2a*) at 24 hpf. *pax2a* is expressed during early somitogenesis in the developing pronephric tubules and is important in establishing the boundary between podocytes and the neck segment of the nephron ^27,28^. In *nup133* morphants, expression of *pax2a* at 24 hpf was similar to that of wild type embryos, suggesting that the tubular development is not compromised in the morphants (Fig. 3a). We further examined the pronephric tubules and ducts in *nup133* knockdown embryos by checking the expression of the kidney specific marker *cadherin 17* (*cdh17*) at 24 hpf and 72 hpf. Again, no obvious difference in expression of *cdh17* was observed between control and *nup133* morphant larvae at 24 or 72 hpf (Fig. 3b). Based on these markers, we conclude that tubular development is not impaired upon *nup133* knockdown.

**Figure 3.**
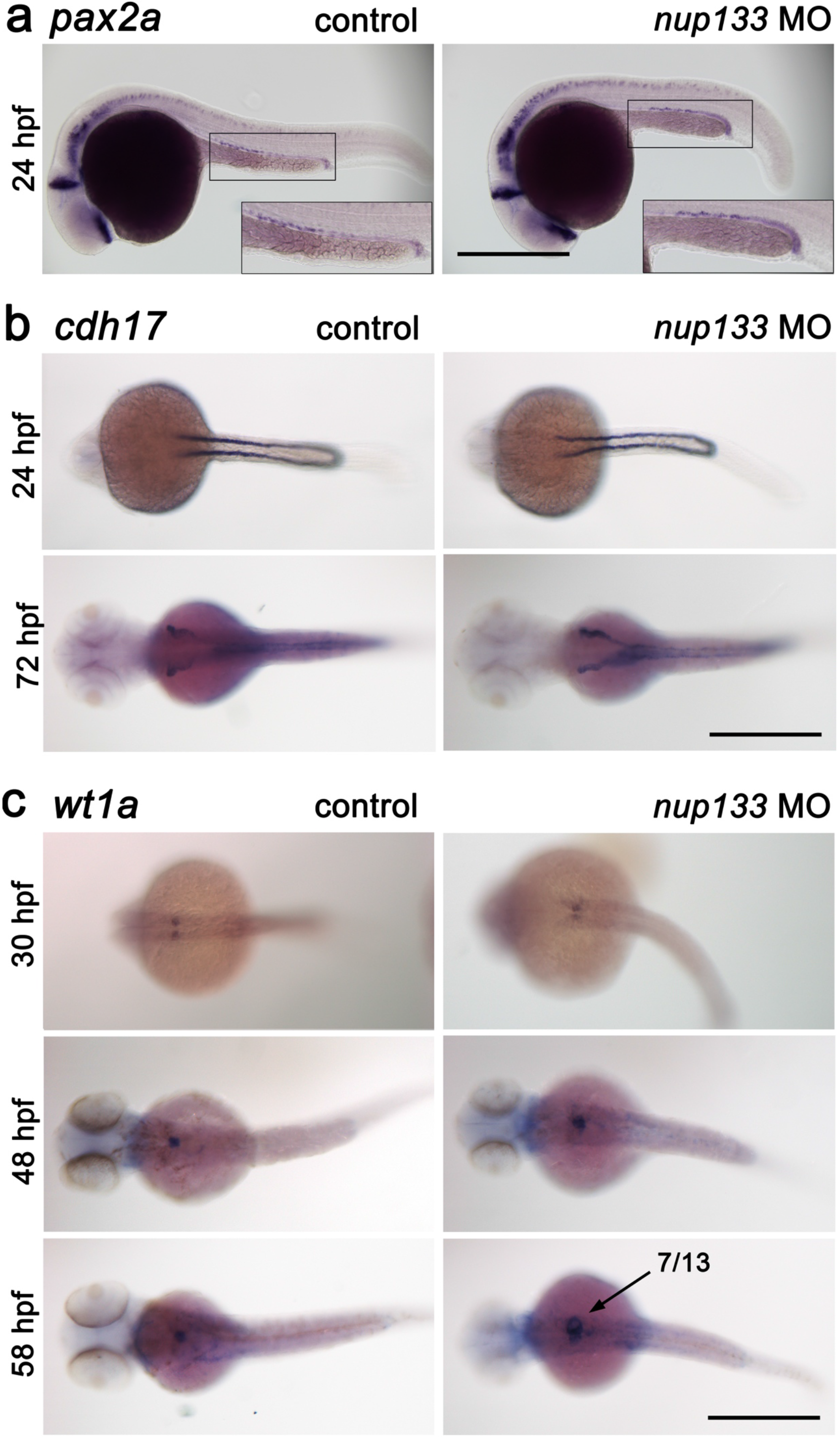
Normal development of the pronephric tubules and glomerulus in *nup133* morphants. (**A**) Lateral view of *pax2a* mRNA expression in pronephric tubules at 24 hpf in uninjected (control) and *nup133* MO-injected embryos reveal that the developmental expression of *pax2a* is not altered in *nup133* morphants. Two-fold magnification of the pronephric tubules is also shown. (**B**) Pronephros marker *cdh17* mRNA expression in *nup133* MO is comparable to that of uninjected controls at 24 and 72 hpf (dorsal view). (**C**) Glomerular development in control and *nup133* MO embryos visualized using the podocyte differentiation marker *wt1a* (dorsal view). At 30 hpf (upper panels), *wt1a* marks future podocytes with two distinct domains in both control and *nup133* MO. At 48 hpf (middle panels), the glomerular primordia merge to the midline to form a single glomeruli. At these stages, mRNA expression does not differ between control and *nup133* MO (middle panels). At 58 hpf (lower panels), after the onset of glomerular filtration, the increased area labeled by the *wt1a* probe (arrow) reflects the glomerular expansion observed in 7 out of 13 *nup133* morphants analyzed. Scale bars, 500 μm.

We next used an mRNA probe for *Wilms tumor suppressor 1a* (*wt1a*) that is predominantly expressed in podocytes throughout pronephric development ^29^. At 24 hpf, *wt1a* is still expressed in two distinct domains, the glomerular primordia, that fuse to form a single glomerulus by 40 hpf ^30,31^. In *nup133* morphants, expression of *wt1a* was comparable at 30 and 48 hpf to that of wild type embryos (Fig. 3c). *wt1a* expression sometimes revealed some glomerular enlargement already at 48hpf, a phenotype that became more evident at 58 hpf, after the onset of the glomerular filtration (Fig. 3c, bottom panels). Nevertheless, *wt1a* transcripts were still expressed in *nup133* morphants podocytes. These results suggest that knockdown of *nup133* does not affect the gross development of zebrafish glomerulus.

### *nup133* deficiency affects the normal function of the glomerular filtration barrier

Because the glomerular enlargement upon *nup133* knockdown becomes striking at the onset of glomerular filtration, we next analyzed the functionality of the pronephros in the morphant larvae. The glomerular filtration barrier (GFB) of the kidney allows the free filtration of water and small solutes, while restricting the flow of large plasma proteins such as albumin ^32^. However, if the GFB is damaged, albumin and other large proteins pass the barrier to a great extent. Part of the albumin is endocytosed by the epithelial cells of the proximal tubules, while the rest gets then excreted via the final urine leading to proteinuria. In zebrafish larvae, it is possible to inject fluorescent compounds of specific size into the general circulation and then to monitor the appearance of fluorescent endosomes in the apical cytoplasm of pronephric tubule cells ^33,34^. To assay for kidney function, we therefore injected fluorescently labeled albumin from Bovine Serum (BSA, Alexa Fluor™ 647 conjugate) into the common cardinal vein of wild type and *nup133* morphant larvae at 4 dpf and then monitored the appearance of fluorescent endosomes in the cytoplasm of pronephric tubule cells. Larvae were fixed 20 minutes after BSA injections and sections of the pronephric proximal tubules were imaged. In the control larvae (n= 6, arising from two distinct experiments), we did not observe fluorescent signal from the injected BSA in the apical endosomes of the proximal tubules, indicating that BSA was not able to pass through the intact filtration barrier (Fig. 4, left panels). In most (5/6 imaged larvae) *nup133* morphants, however, fluorescently labeled endosomes were detectable in the proximal tubules (Fig. 4, right panels). This suggests that the filtration barrier of the glomerulus is impaired upon *nup133* knockdown, allowing the passage of macromolecules with a size exceeding the size selectivity of an intact GFB, a condition defined in patients as proteinuria.

**Figure 4.**
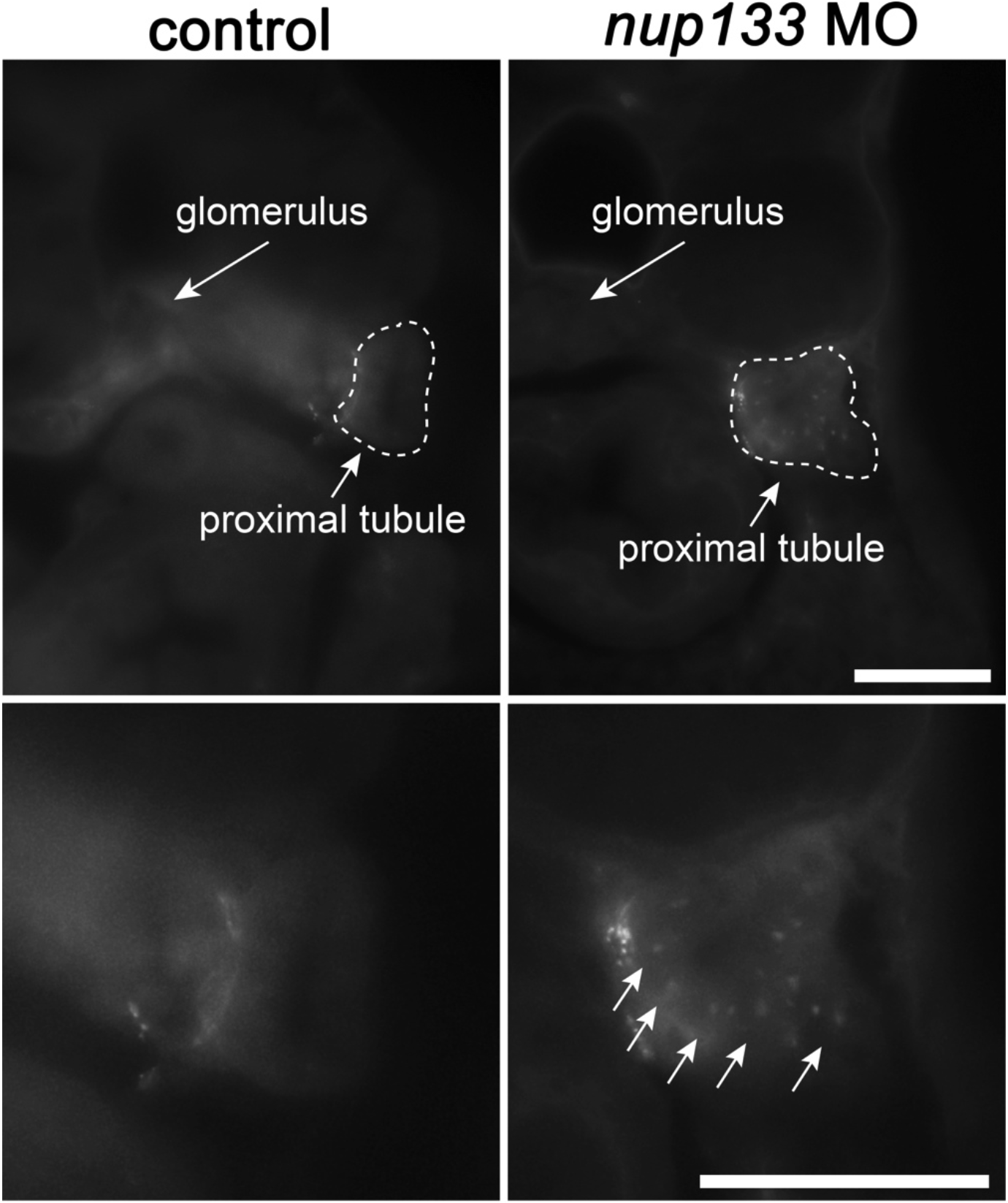
Glomerular filtration is impaired in *nup133* morphants. Representative images of cross sections of 4 dpf control (left panels) and *nup133* morphants larvae (right panels) fixed 20 minutes after injection of Alexa Fluor™ 647 conjugated-BSA into the common cardinal vein. Lower panels represent a higher magnification view of the proximal tubule region (dotted lines). Note the uptake of fluorescent BSA in the apical endosomes of the proximal tubules of the Nup133-depleted larva (arrows). Scale bars 50 μm.

### *nup133* knockdown results in podocyte FPs effacement

In order to determine whether the glomerular filtration impairment observed in *nup133* morphants was a consequence of a defect in the glomerular filtration barrier (GFB), we conducted electron microscopy studies to compare the ultrastructure of the glomerulus in *nup133* morphants and wild type larvae at 5 dpf, a time point when the larval glomerulus development is considered nearly complete ^30,35^. In wild type larvae, the glomerulus showed the typical organization with podocytes harboring well-developed interdigitated foot processes on the outer side, and fenestrated endothelial cells on the inner side of the GBM (Fig. 5b, left panels). In *nup133* knockdown larvae, however, a moderate foot process effacement was observed in many areas of the glomerulus, with no detectable alteration of the fenestrated endothelial cells (Fig. 5b, right panels). These data thus indicate that zebrafish *nup133* contributes to the structural integrity of the GFB.

**Figure 5.**
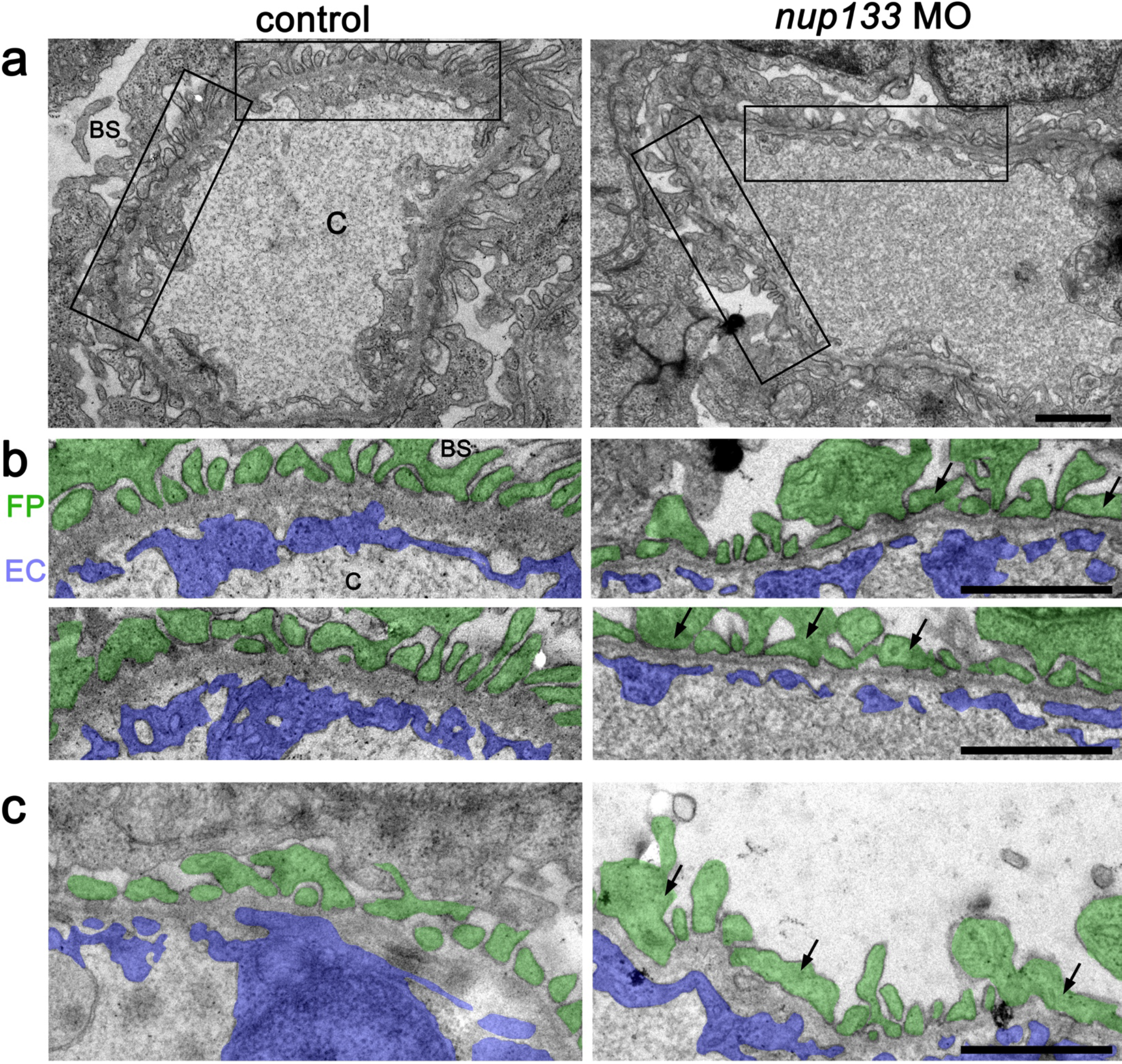
*nup133* knock down induces moderate foot processes effacement. (**A**) Transmission electron micrographs showing the glomerular capillary wall of 5 dpf uninjected control (left panels) and *nup133* MO injected larvae (right panels). (**B**) Two-fold magnification of the areas indicated in (**A**), and (**C**) representative images from distinct larvae were pseudocolored to better highlight the glomerular filtration barrier components (podocyte foot processes in green, fenestrated endothelium in blue). Abbreviations: BS: bowman space; C: capillary lumen; EC: endothelial cells; FP: foot processes Arrowheads point to irregular-shaped foot processes that are more frequently found in *nup133* MO larvae. Scale bars 1 μm.

## Discussion

In this study we used zebrafish as model organism to evaluate the contribution of Nup133 to vertebrate kidney function. Using MO-based depletion of Nup133 followed by rescue experiments with *nup133* mRNA, we have identified *nup133* as a new regulator of glomerular structure and functional integrity.

While we report here a kidney-specific alteration caused by Nup133 depletion in zebrafish, more severe developmental phenotypes were previously observed upon *Nup133* inactivation in mouse ^19^. This likely reflects the complete absence of Nup133 protein in the mouse mutant as compared to its incomplete MO-mediated depletion in zebrafish. Of note, the extent of depletion of structural nucleoporins may also lead to distinct phenotypes as revealed by previous studies of zebrafish *nup107*. Indeed, while MO-induced depletion of zebrafish Nup107 was reported to cause glomerular defects including podocytes foot processes effacement in one study ^16^, Nup107 deficiency caused by transposon insertion in *nup107*^*tsu068Gt*^ transgenic embryos was reported to affects multiple tissues including the pharyngeal skeleton, the intestine, the swim bladder and the eyes ^36,37^.

Our study of glomerulus and proximal tubules expression markers shows that the decreased level of Nup133 does not affect the initial stages of nephron development. In addition, TEM analysis of the glomerulus of Nup133-depleted zebrafish reveals that all the cell types constituting the GFB, including the podocytes, are in place. However, the podocytes of Nup133-depleted zebrafish present some degrees of effacement that correlate with a defective GFB. This result therefore suggests that Nup133 plays a fundamental role in the maintenance rather than in the establishment of the refined podocytes structure required for GFB function.

Several mechanisms could explain the implication of Nup133, and more generally, of Y-complex Nups, in glomerular function and thus NS. Nup133 deficiency may specifically alter the nuclear transport of signaling molecules, as previously reported for Nup93 that belongs to another structural domain of the NPC ^14^ Indeed, human podocytes and HEK293 cells expressing Nup93 mutations identified in SRNS patients exhibited impaired nuclear import of SMAD4 upon stimulation with bone morphogenetic protein 7 (BMP7), a secreted molecule involved in kidney development and response to renal injury. This result was consistent with a previous study in which depletion of Drosophila Nup93 led to a defective import of the SMAD4 orthologue ^38^. Noteworthy, the latter study also revealed the implication of two Drosophila Y-Nups, Nup85/75 and Sec13, but not of Nup133, in this specific import pathway ^38^. While we cannot formally exclude Nup133 contribution to the nuclear import of SMAD4 in vertebrates, Nup133 depletion may also affect other signaling cascades required for the maintenance of an intact GFB. Indeed, two Y-complex subunits, Nup107 and Nup37, were reported to be regulators of the ERK and YAP pathways, respectively ^39,40^.

Nup133 knockdown may also alter nuclear mechanotransduction, for instance by interfering with the ‘linker of nucleoskeleton and cytoskeleton’ (LINC) complex that establish a stable and cross-linked network between the cytoskeleton and the lamina underneath the NE (reviewed in ^41^). Indeed, while Nup133 is required for the proper assembly of the NPC basket ^42^, intimate links between the NPC basket, the LINC subunit Sun1 and Lamin-C have been reported ^43,44^ As previously discussed ^16^, deregulation of mechanotransduction signaling pathways may in turn affect podocytes that are subjected to mechanical stretching caused by capillary pressure. As reported for a subset of nucleoporins (reviewed in ^45^), Nup133 may be involved in gene regulation. It may thus modulate the expression of specific genes required for proper function of the GFB, notably podocytes. Finally, one may also keep in mind that several nucleoporins localize at the base of the cilia ^46,47^. In particular, another member of the Y-complex, Nup85, was recently reported to be required for proper cilia localization of Nup98, a nucleoporin that regulates diffusion of soluble molecules through the ciliary base ^48^. Moreover, mutations affecting the kinetochore protein Cenp-F, an established partner of Nup133 ^49^ were identified in patients affected by severe ciliopathy and microcephaly ^50^. Since appearance of kidney cysts in zebrafish can also be caused by inactivation of several ciliopathy genes (reviewed in ^51,52^), the hypothesis that Nup133 depletion may contribute to kidney cysts formation in zebrafish by affecting cilia, might deserve attention in the future.

So far, the only Y-complex nucleoporin gene found to be mutated in SRNS is *Nup107* ^15–18^. Our data now indicate that Nup133, its direct partner within the Y-complex, is also required for proper glomerular function in zebrafish. In addition, a recent study revealed that normal expression level of another Y-complex component, Nup160, is critical for the proliferation and viability of podocytes *in vitro* ^53^. These data indicate that in addition to *Nup107*, at least these two other Y-Nups, or possibly all the nine Y-complex subunits (*Nup107, Nup133, Nup160, Nup96, Nup85/75, Nup43, Nup37, Sec1,* and *Sec13*) are candidate genes that would be worth testing in SRNS patients for which the mutated gene is yet unidentified. Our study, together with an increasing number of studies correlating single-gene mutations in *Nups* with cell type-specific defects, thus contributes to strengthen the rise of the “nucleoporopathies” concept ^1^ an exciting development in the nuclear pore complex field.

## Materials and methods

### Fish maintenance and breeding

Zebrafish (*Danio rerio*) were kept at 26°C under a 14-h/10-h light/dark cycle and bred as previously described ^54^. Larval stages were raised at 28°C in E3 embryo medium (5 mM NaCl, 0.17 mM KCl, 0.33 mM CaCl_2_, 0.33 mM MgSO_4_) containing 0.01% methylene blue. 0.003% PTU (1-phenyl-2-thiourea; Sigma Alrdrich) was added in embryo medium to inhibit melanin synthesis during larval development and facilitate fluorescent microscopy. *Tg(wt1b:EGFP)* line was a kind gift from Dr. Christoph Englert (Leibniz Age Research, Jena, Germany). All experiments were performed in accordance with internationally recognized and with Swiss legal ethical guidelines for the use of fish in biomedical research and experiments were approved by the local authorities (Veterinäramt Zürich Tierhaltungsnummer 150), the European Council Directive for animal use in science (2010/63/EU) and were approved by the local authorities (Veterinäramt Zürich TV4206).

### Whole-mount in situ hybridization

Sequences were identified and annotated using combined information from expressed sequence tags and genome databases (GeneBank, http://www.ncbi.nlm.nih.gov; Ensembl, http://www.ensembl.org/index.html). The primers used for probe preparation are listed in supplementary Table S1. *nup133, pax2a, wt1a* and *cdh17* cDNAs were all isolated by RT-PCR from total RNA from 24-48 hpf embryos and cloned into TOPO pCRII vector (TA Cloning Kit Dual Promoter, Invitrogen) as previously described ^55^. The resulting plasmids were linearized for SP6 and T7 *in vitro* transcription and purified with phenol-chloroform. Digoxigenin (DIG)-labeled antisense (or sense, as negative control for *nup133* expression) RNA probes were generated using DIG-RNA-labeling kit (Roche Diagnostic). Zebrafish embryos were fixed in 4% paraformaldehyde in phosphate-buffered saline (PBS) at 4°C overnight and whole mount in situ hybridization was performed as previously described ^56^. Briefly, on day 1 the larvae were treated with proteinase K and then fixed with 4% paraformaldehyde (PFA) before prehybridization at 64°C. Hybridization of RNA probes was done at 64°C overnight. On day 2, after several stringency washes at 64°C, probes were blocked in 1× Roche blocking solution in Tris/NaCl/Tween. Anti-DIG AP antibody was applied overnight at 4°C. On day 3, after several washing steps, signal was detected by incubation in staining buffer. Stained embryos were postfixed with PFA and imaged in glycerol with an Olympus BX61 light microscope or with a stereomicroscope (Olympus MVX10). Images were processed and assembled using Adobe Photoshop and Adobe Illustrator CS6.

### Cryosectioning

Embryos at desired stages were fixed in 4% PFA at 4°C overnight, cryoprotected in 30% sucrose overnight and embedded in tissue freezing medium (Richard-Allan Scientific Neg-50 Frozen Section Medium Thermo Fisher Scientific). 14-16 μm transverse sections were cut on a Microm HM 550 cryostat and imaged with a Olympus BX61 wide-field microscope (Volketswil, Switzerland).

### EdU incorporation and detection

5 dpf zebrafish larvae were incubated in E3 medium containing 10 mM EdU in 1 % DMSO for 1h in the dark, followed by 1h recovery time in E3 medium. Larvae were fixed in 4% PFA for 1h, rinsed in PBT and processed for cryosections. EdU staining was then performed as previously described ^57^.

### Morpholino and RNA injections

The *nup133* gene was targeted with specific antisense splice-blocking Morpholinos (MOs) (GeneTools, Philamath, OR) designed to target the exon-intron boundary (splice donor, *nup133*_E3I3) of exon 3 and the intron-exon boundary (splice acceptor, *nup133*_I3E4) of exon 4, respectively (MO sequences are provided in Supplemental Table S1). The morpholinos were diluted in RNase-free water with 0.1% phenol red as an injection tracer and ~1 nl was injected into fertilized embryos at 1-2 cell stage. Uninjected sibling embryos and embryos injected with standard control (SC) morpholinos were used as controls. Evaluation of morphological changes was performed at 3 dpf. A range of concentration was first used to determine the optimal MO amount required to induce a specific phenotype without inducing toxicity. The two MOs were then always injected at 3 mM each. To examine splicing defects caused by *nup133*_E3I3 and *nup133*_I3E4 injection, poly(A) RNA were extracted from control or MO-injected embryos at 1 dpf and RT-PCR and characterized by RT-PCR using primers designed from flanking exon-coding sequence (Figure 2a and Supplementary Table S1 online) The RT-PCR products were purified from gel, re-amplified with nested primers and altered splicing was confirmed by sequencing.

For the rescue experiments, a pBluescript KSM vector containing *3xHA-mCherry-Dr nup133* was used. The plasmid was generated using PCR amplification using proofreading DNA polymerases (Phusion HF, NEB) and NEBuilder HiFi DNA Assembly Cloning Kits. Briefly, the sequence encoding three copies of the influenza virus *hemagglutinin* (*HA*) epitope was amplified from pFA6a-3HA-kanMX6 ^58^, the *mCherry* were amplified from plasmid #1937 ^42^, full-length zebrafish *nup133* was amplified from a Dharmacon cDNA vector (Clone ID: 2600558) and the three PCR products were inserted by recombination in a pBluescript KSM vector (Stratagene). PCR-amplified fragments and junctions were checked by sequencing. Plasmid map is available upon request. For *in vitro* mRNA production the *3xHA-mCherry-Dr nup133* coding fragment was linearized by digestion with the restriction enzyme NotI and purified with phenol-chloroform extraction. Capped and tailed RNA was *in vitro* transcribed using the mMessage mMachine T3 kit (Life Technologies, Zug, Switzerland) according to the manufacturer’s instruction. A polyA tail was added with polyA tailing kit (Invitrogen by thermos Fischer Scientific) followed by purification with MEGAclear Transcription Clean-Up Kit (Ambion). The mRNA (300nM) was injected into the embryos at one cell stage before MO injection.

### Whole mount live-animal imaging

To evaluate morphological changes, larvae were anesthetized with 200 mg/ml 3-aminobenzoic acid methyl ester (MESAB, Sigma-Aldrich), mounted in 1.5% low melting temperature agarose in E3 medium and imaged using a stereomicroscope (Olympus MVX10) equipped with a color camera (ColorViewIII, Soft imaging System, Olympus).

### Western blotting

For western blot analysis of 3xHA-mCherry-Dr Nup133 expression, groups of 20-40 embryos at 24 hpf were homogenized by sonication in 50-100 μl of Laemmli buffer. The lysates were separated on NuPAGE™ 4–12% Bis-Tris gels (using MOPS as running buffer) and transferred to nitrocellulose membranes. The resulting blots were stained using Ponceau, saturated with TBS, 0.1% Tween, and 5% dried milk, and probed with mouse monoclonal antibody HA.11 (clone 16B12; Eurogentec #MMS-101R; 1:2,000). Incubations of the membrane with primary and HRP-conjugated secondary antibodies (Jackson ImmunoResearch Laboratories) were done in TBS buffer (0.1% Tween, 5% dried milk), and signals were detected by enhanced chemiluminescence (SuperSignal^®^ Femto; Thermo Scientific).

### Transmission electron microscopy and histology

Zebrafish larvae at 5 dpf were fixed in 2.5% glutaraldehyde in 0.1 M cacodylate buffer, pH 7.2 overnight. To achieve a better penetration of the fixative, the tail of each larva was cut off with a scalpel prior to fixation. The larvae were rinsed in 0.1 M cacodylate buffer before postfixation in 1% osmium tetroxide in cacodylate buffer for 1 hour at room temperature. The samples were rinsed in distilled water before en block staining in 1% aqueous uranyl acetate for 1 hour at room temperature. Following dehydration though a graded series of ethanol ranging from 50% to 100%, the larvae were infiltrated overnight in 66% Epon/Araldite in propylene oxide. Finally, the specimens were embedded in 100% Epon/Araldite and placed in a polymerizing oven at 60 °C for 26 h. Semi thin section (2 μm) were stained with toluidine blue and used for histological studies. Ultrathin sections (65 nm) of the glomerulus and the proximal tubules, obtained using a Leica EM FCS ultramicrotome were collected on formvar coated copper grids, stained in lead for 5 minutes to increase the contrast, and examined with a Philips CM-100 scope at 80 kV. Images were acquired using the Gatan Microscopy Software.

### Glomerular filtration assay

Zebrafish larvae at 4 dpf were anesthetized with 200 mg/ml 3-aminobenzoic acid methyl ester (MESAB, Sigma-Aldrich). 1 nl of Albumin from bovine serum (BSA), Alexa Fluor 647 conjugate (Thermoischer scientific) was injected into the common cardinal vein according to ^34^. The larvae were transferred to E3 medium for recovery. 20 mins after injections, the larvae were fixed in 4% PFA overnight and processed for cryosections and imaging.

## Acknowledgements

We are grateful to Dr. Nathan W. Luedtke (Chemistry Department, University of Zurich) for kind help with EdU staining and to Dr. Christoph Englert (Leibniz Age Research, Jena, Germany) for sharing the *Tg(wt1b:EGFP)* line. We also acknowledge Zhiyong Chen, Yuya Sugano, Ruxandra Bachmann-Gagescu and Matthias Gesemann for support and helpful advices, Benoit Palancade for critical reading of the manuscript and other members of our labs for helpful advices. The authors acknowledge the assistance and support of the Center for Microscopy and Image Analysis, University of Zurich for support with electron microscopy experiments. We also acknowledge the ImagoSeine core facility of the Institut Jacques Monod, member of IBISA and of the France-Bioimaging (ANR-10-INBS-04) infrastructures. Work in the laboratory of VD is supported by the Centre National de la Recherche Scientifique (CNRS), the “Fondation pour la Recherche Médicale” (Foundation for Medical Research) under grant No DEQ20150734355, “Equipe FRM 2015” and the Labex Who Am I? (ANR-11-LABX-0071; Idex ANR-11-IDEX-0005-02). AB received PhD fellowships from the “Ministère de l’Enseignement Supérieur et de la Recherche” and the “Ligue Nationale contre le Cancer” and a “transition post-doc” grant from the Labex Who Am I?. Work in the laboratory of SN was supported by project grants from the Swiss National Science foundation (to SCFN and JL) and the RiMED foundation in Palermo (to CCC). CCC received a fellowship from the RiMED foundation.

## Author Contributions

CCC, AB, JL, SN and VD conceived and designed the experiments; CCC, AB and MH performed the experiments; CCC, AB, JL, SN and VD analyzed the data; CCC, AB and VD wrote the manuscript with contribution from all co-authors.

